# IN SILICO DESIGN AND BINDING MECHANISM OF E3 LIGASE UBR1 RECRUITERS

**DOI:** 10.1101/2024.01.14.575594

**Authors:** Miguel A. Maria-Solano, Raudah Lazim, Sun Choi

**Affiliations:** Global AI Drug Discovery Center, College of Pharmacy and Graduate School of Pharmaceutical Science, Ewha Womans University, 03760 Seoul, Republic of Korea

## Abstract

Proteolysis Targeting Chimeric Molecules (PROTACs) represent a promising avenue in drug discovery, as they can induce the targeted degradation of disease-relevant proteins within the cellular machinery. These compounds comprise a ligand tailored to bind the specific targeted protein connected to a recruiter molecule that engages with the E3 ligase. Despite their promise as therapeutic agents, their clinical advancement has encountered substantial challenges, primarily due to the limited availability of suitable E3 ligases. Additionally, cell permeability and proteolytic stability due to their peptide nature often hamper their application. In this study, we focus on the development of recruiters for the E3 ligase UBR1. This widely expressed protein has recently been demonstrated to be efficient in driving the degradation of oncogenic proteins. Our computational approach leverages a fragment-based peptidomimetics strategy, integrating pharmacophore filtering, docking, and fragment-linking optimization. Finally, we subject the wild-type peptide and the most promising combined fragments to advanced binding free energy calculations, unveiling insights into their dynamic water-mediated binding mechanisms and their potential as robust E3 ligase UBR1 recruiters, ultimately leading to the identification of promising compounds. This computational workflow is readily applicable to the development of related PROTACs and also to model protein-protein interactions with similar characteristics.

## INTRODUCTION

Proteolysis Targeting Chimeric Molecules (PROTACs) are heterobifunctional molecules that induce the approach of a target protein to the cellular degradation machine.[1-3] They consist of three essential components: a ligand that binds to the target protein, a recruiter that binds to the E3 ligase, and a linker that connects these two elements. When introduced into the cellular environment, PROTACs form a ternary complex involving the target protein and the E3 ligase. This complex triggers the polyubiquitination of the target protein, marking it for subsequent degradation by the proteasome (**Figure 1**).

**Figure 1.**
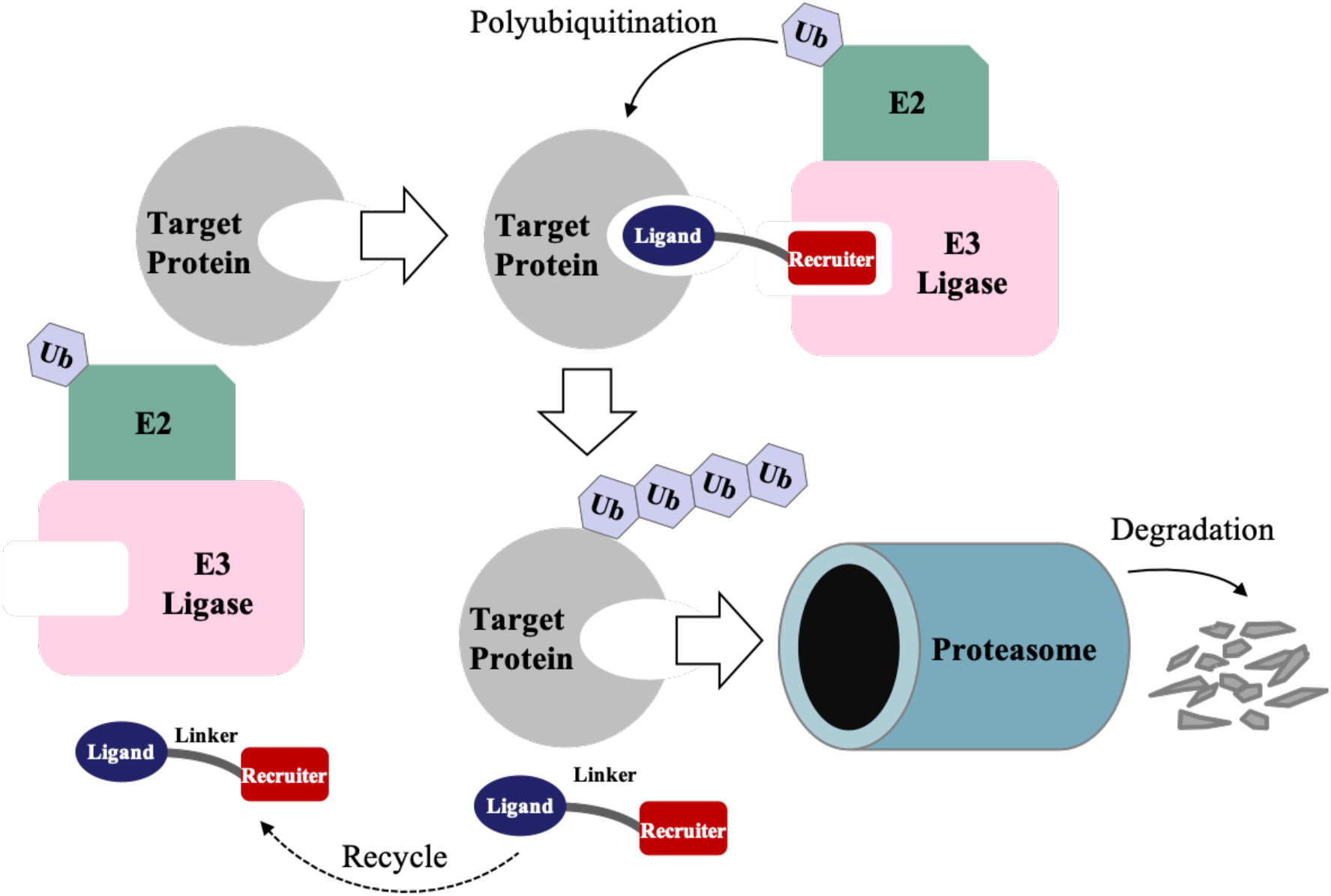
Overview of Proteolysis Targeting Chimeric Molecules (PROTACs) induced-degradation mechanism. PROTACs consisting of a protein-target ligand and an E3 ligase recruiter connected by a linker induce the polyubiquitination and subsequent proteasomal degradation of target proteins.

The degradation of disease-relevant proteins is an emerging therapeutic strategy for a wide range of diseases, such as cancer, viral infections, and immune and neurodegenerative disorders.[4-6] PROTACs have demonstrated significant advantages over classic inhibitors, particularly in selectivity and their ability to overcome drug resistance issues. This can be attributed to their induced-degradation mode of action. Once the target protein is marked for proteasomal degradation, PROTACs can dissociate and be recycled for subsequent degradation events. Therefore, PROTACs do not rely on high binding affinities to exhibit potent degradation activities. In contrast, classical small-molecule inhibitors operate through competitive and occupancy-based mechanisms, requiring deep, well-defined binding pockets and high binding affinities. This makes classical inhibitors more susceptible to resistant mutations and target expression increases.[4, 7] The initial step in designing a PROTAC involves the identification of the target protein and an appropriate E3 ligase. PROTACs can be designed to degrade known protein targets by exploiting existing inhibitors as ligands,[8] but they also offer the potential to explore the “undruggable” proteome that is not susceptible to classic inhibitors.[9] However, the identification of an efficient cellular degradation machinery is more problematic.[10] Despite the existence of approximately 600 estimated E3 ligases members,[11] only a few have been utilized, with cereblon (CRBN) and Von Hippel-Lindau (VHL) being the most commonly employed E3 ligases.[12, 13] The limited repertory of available cellular degradation machines hampers the development of PROTACS, as they are unable to selectively degrade every protein target, and the expression levels of these E3 ligases are insufficient in certain cell types.[10] In this context, the N-Degron pathway,[14, 15] which is a proteolytic system recognizing N-terminal residues of short-lived proteins through the E3 ligase UBR family, has been recently investigated.[16] Particularly, UBR proteins are ubiquitously expressed in most cells.[17] Consequently, PROTACs containing a recruiter for E3 ligase UBR could be exploited to degrade target proteins regardless of the cell type.[18] In this regard, a rationally designed PROTAC consisting of an N-Degron peptide as recruiter linked to a staple peptide as ligand successfully degraded the steroid receptor coactivator-1 (SRC-1), recognized as an oncogenic protein. This study highlights the potential of the N-Degron pathway for cancer treatment.[16, 19] However, the clinical progression of PROTACs has encountered obstacles due to their poor cell permeability and proteolytic stability, which is often attributed to their peptide nature and large molecular weight. Furthermore, the large and shallow protein-protein interaction (PPI) surfaces[20-22] make the design of small molecules with efficient E3 ligase recruiting activity a difficult task.

To overcome such obstacles, we employed an *in silico* peptidomimetics strategy, which involved the virtual screening of compound libraries to identify small molecules that mimic the structural features of N-Degrons responsible for its binding affinity while providing higher stability towards proteolysis, better transport properties, and selectivity.[20, 23] Hence, our focus was on designing E3 ligase UBR1 recruiters, aiming to identify a potential set of compounds that could be linked to protein target ligands and effectively degrade disease-relevant proteins in any cell type. Specifically, we developed a fragment-based computational strategy that combined pharmacophore filtering, docking, and fragment-linking optimization. Subsequently, the resulting combined fragments were filtered as a function of docking score, drug-likeness, and synthetic accessibility. Finally, the combined fragments exhibiting the best scores were further evaluated through advanced free energy calculations, which revealed the molecular basis of the binding mechanism and the potential of these designed compounds as effective E3 ligase UBR1 recruiters.

## RESULTS AND DISCUSSION

### Fragment pharmacophore-based virtual screening of the N-Degron-UBR box comple

Structure-based virtual screening (SBVS) is a well-established and successful computational methodology that exploits the three-dimensional knowledge of the protein target to select drug candidates from large compound databases.[24] Fortunately, crystal structures of N-Degron peptides in complexes with UBR1 have been described. [25] N-Degron peptides bind in a relatively shallow acidic cleft located at the UBR box domain. Structural inspection and mutagenesis experiments indicated that the first residue (positively charged) of N-Degron plays a major contribution to the binding affinity while the second residue (hydrophobic) plays a minor role. This is evidenced by the co-crystalized structure of the wild-type peptide from cohesion subunit Scc1 (RLGES), where the guanidinium group and NH3+ groups of the first residue (Arginine) establish an extensive hydrogen bond network with I174, D176, and D179 residues (Figure 2A). Considering this structural data (PDB 3NIN),[25] we generated a pharmacophore model, which mostly contained positive ionizable areas and hydrogen bond donors. It also contained a hydrophobic area and some hydrogen bond acceptors (**Figure 2B**). Given that the N-Degron and UBR box interface involves protein-protein interactions (PPIs), we started screening PPI commercial libraries. However, there were no compounds that matched the physicochemical properties of the pharmacophore model generated based on the N-Degron/UBR box interface. This is because this specific case of PPI produces a pharmacophore with challenging physicochemical properties, including a long length (ca. 15 Å) that is hard to target. To overcome this challenge, we decided to develop a fragment-based pharmacophore model.[26] Specifically, we divided the original pharmacophore into three zones giving rise to three fragment pharmacophore models: FP-A (zone 2+zone 3), FP-B (zone 1+zone 2), and FP-C (zone 3), see **Figure 2C**. Subsequently, we performed the screening of fragment libraries using the three fragment pharmacophore models to find compounds that bind to each zone. Fragment libraries are composed of small libraries of low-complexity compounds, which have been proven to present high hit rates. In total, considering the screening of all commercial libraries, we obtained 1,511 fragment hits, from which 97 were identified in the FP-A screening, 480 in the FP-B, and 932 in the FP-C (**Supplementary Table 1**). An interesting aspect is that FP-A presents the lowest hit rate. This indicates that zone 3 is more difficult to target than zone 1. However, as described previously, zone 1 is the most critical to the binding energy. The resulting fragment hits are expected to be complementary to the UBR box binding site at their respective binding zones. Nevertheless, to obtain compounds with good binding affinities, these fragments must be further optimized through expansion strategies, such as fragment growing, merging, and linking. This is an attractive approach to discovering new chemical entities with high specificity. [27]

**Figure 2.**
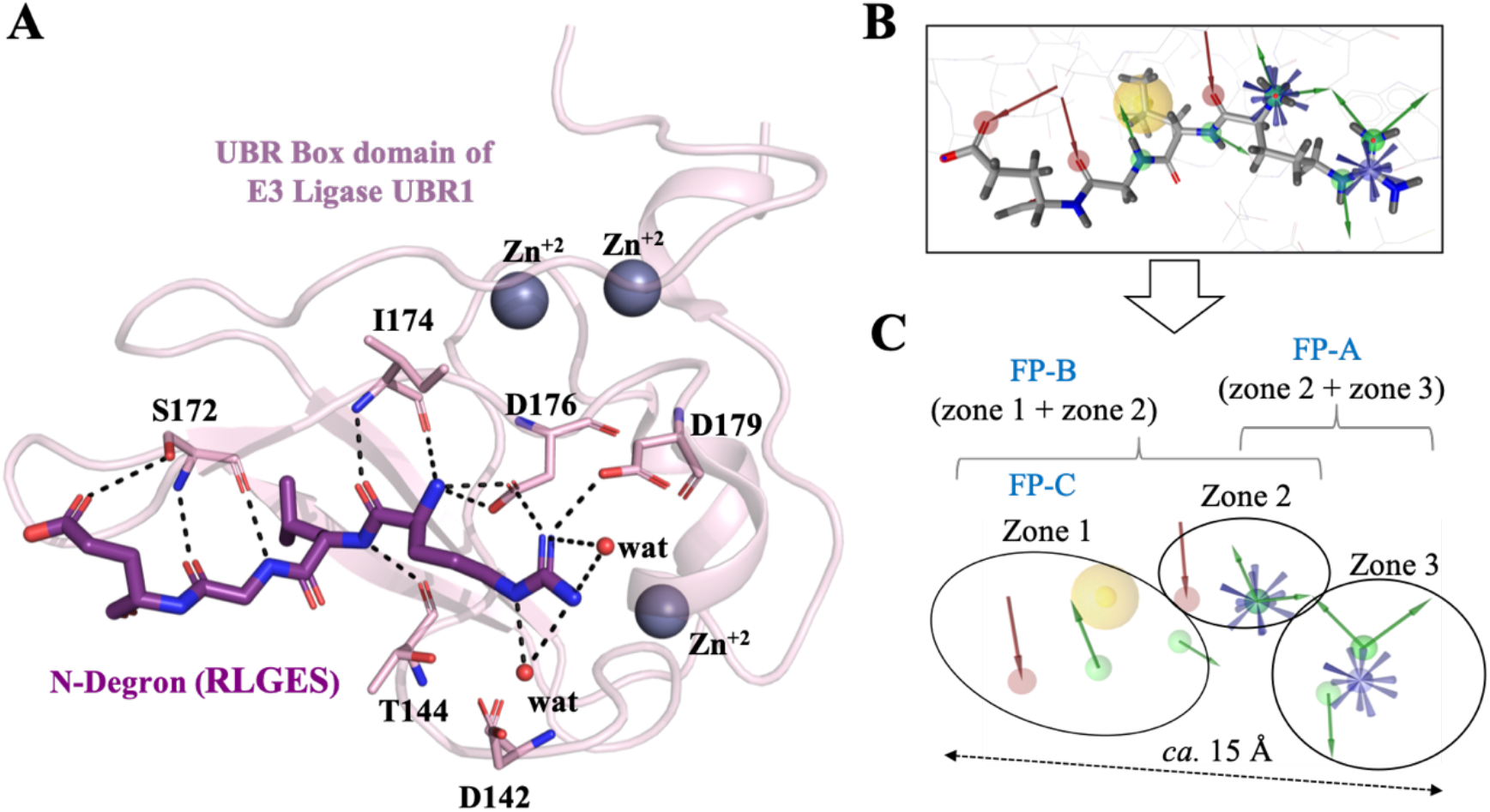
Structural information and fragment-based pharmacophore model. **(A)** View of the RLGES N-Degron peptide in complex with the UBR box domain of E3 Ligase UBR1 (PDB 3NIN). The polar contacts between the RLGES peptide and the UBR box residues are shown as black dashed lines. **(B)** Original pharmacophore model derived from PDB 3NIN data. It is composed of positive ionizable areas (in blue), hydrogen bond donors (in green) hydrogen bond acceptors (in red), and hydrophobic interactions (in yellow). **(C)** Fragment-based pharmacophore model consisting of three fragment pharmacophores (FP-A, FP-B, and FP-C).

### Docking of fragment hits in the UBR box followed by linking optimization

Molecular docking is an efficient methodology to rank a large number of compounds based on the estimation of binding affinity. [28] More importantly, it offers the 3D visualization of the virtual hits docked in the binding pocket. This structural information aids in the execution of hit optimization procedures. In this work, a fragment linking optimization protocol as described in the “combine fragment” panel of Schrodinger software was used. First, the 1,511 fragments hits were docked in the UBR box using Glide Docking and second, the fragments that were docked in proximity were joined by identifying bonds that can be formed according to geometrical criteria. For fragment pairs that have no atoms close enough to create new bonds but still not far apart, at most two methylene linkers can be included, see an example of a combined fragment in **Figure 3A**. Among the combined fragments generated, the top-scored 150 docked molecules were selected for further analysis. Considering the docking score and the ligand efficiency (docking score/number of heavy atoms) of the RLGES peptide as our reference values (docking score: - 12.63 and ligand efficiency: 0.32), we established threshold values of -10 for the docking score and -0.3 for the ligand efficiency, resulting in 12 combined fragments. These results indicate that we successfully generated efficient combined fragments that present similar docking values to those of the RLGES peptide. However, a new cryo-EM structure of E3 ligase UBR1 has been successfully determined, including the complete UBR1 structure in complex with Ubiquitin-conjugating enzyme (ubc2) Ubiquitin (Ub) and N-Degron (PDB 7MEX).[29] Based on this newly solved structure, we considered the potential of the protein surroundings of the UBR box to cause steric effects to the docked combined hits by scrutinizing the structure of the E3 ligase-UBR1 complex and the UBR box domain (PDB 3NIN) used for docking. The overlap between the UBR box domain used in the docking study with the complete E3 ligase UBR1 structure (PDB 7MEX) showed that the docking poses of the combined fragments can extend to regions occupied by UBR1, thus causing steric effects (**Figure 3B**). Based on this observation, we repeated the docking and linking optimization of the fragments using the complete UBR1 structure. Applying the same docking score threshold (vide supra), we obtained a total of 26 combined fragments, all of them exhibiting superior docking score and ligand efficiency compared to the RLGES peptide (**Figure 4**). Note that the RLGES peptide presents a slightly higher docking score when docked to the complete UBR1 structure. Interestingly, the resulting combined fragments are complementary to those obtained using the UBR box domain only but the binding poses obtained formed more favorable interactions with the UBR1 surroundings, yielding a better fit in the UBR1 binding interface (**Figure 3C**). This suggests that the consideration of the protein surroundings of UBR1 during the virtual screening process may be important for the design of efficient E3 ligase UBR1 recruiters.

**Figure 3.**
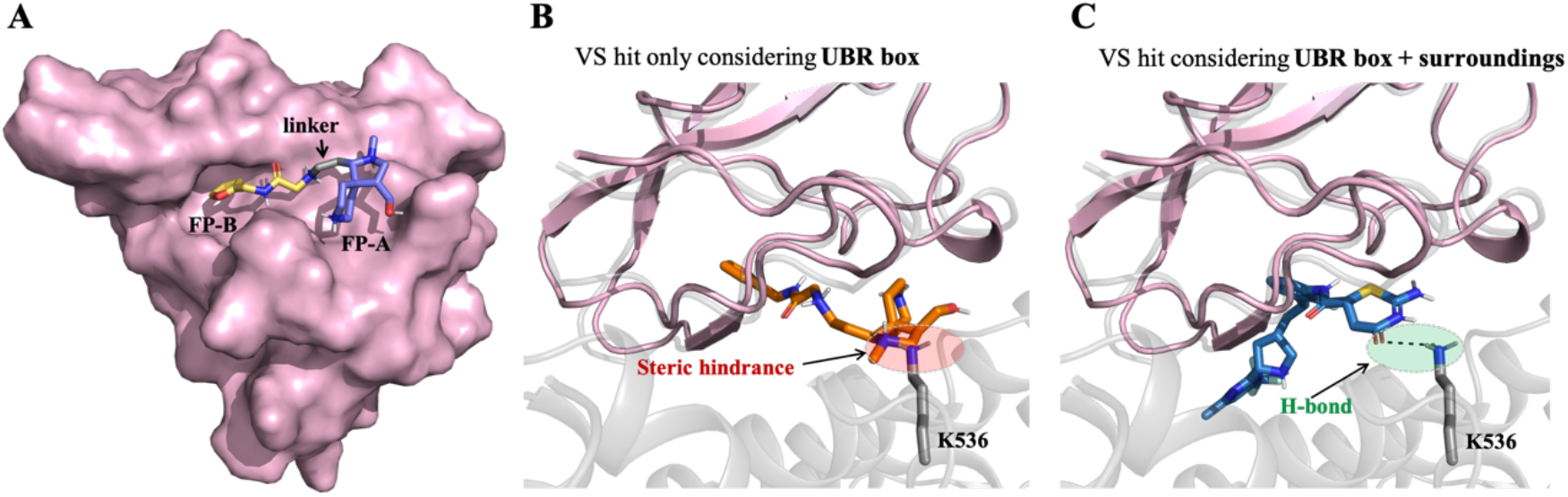
Representative combined fragments docked in UBR1. **(A)** Example of combined fragment docked in the UBR box (shown as a pink surface). The combined fragment is composed of an FP-B fragment (yellow-orange), an FP-A fragment (slate), and an ethylene linker (grey). **(B)** and **(C)** Overlap of the UBR box domain (PDB 3NIN), in pink, and the complete length chain of E3 ligase UBR1 (PDB 7MEX), in grey. The docked combined fragment (orange sticks) only considering the UBR box domain in the docking process performs steric clashes with the UBR box protein surroundings, while the docked combined fragment (blue sticks) considering the complete UBR1 protein performs favorable interactions.

**Figure 4.**
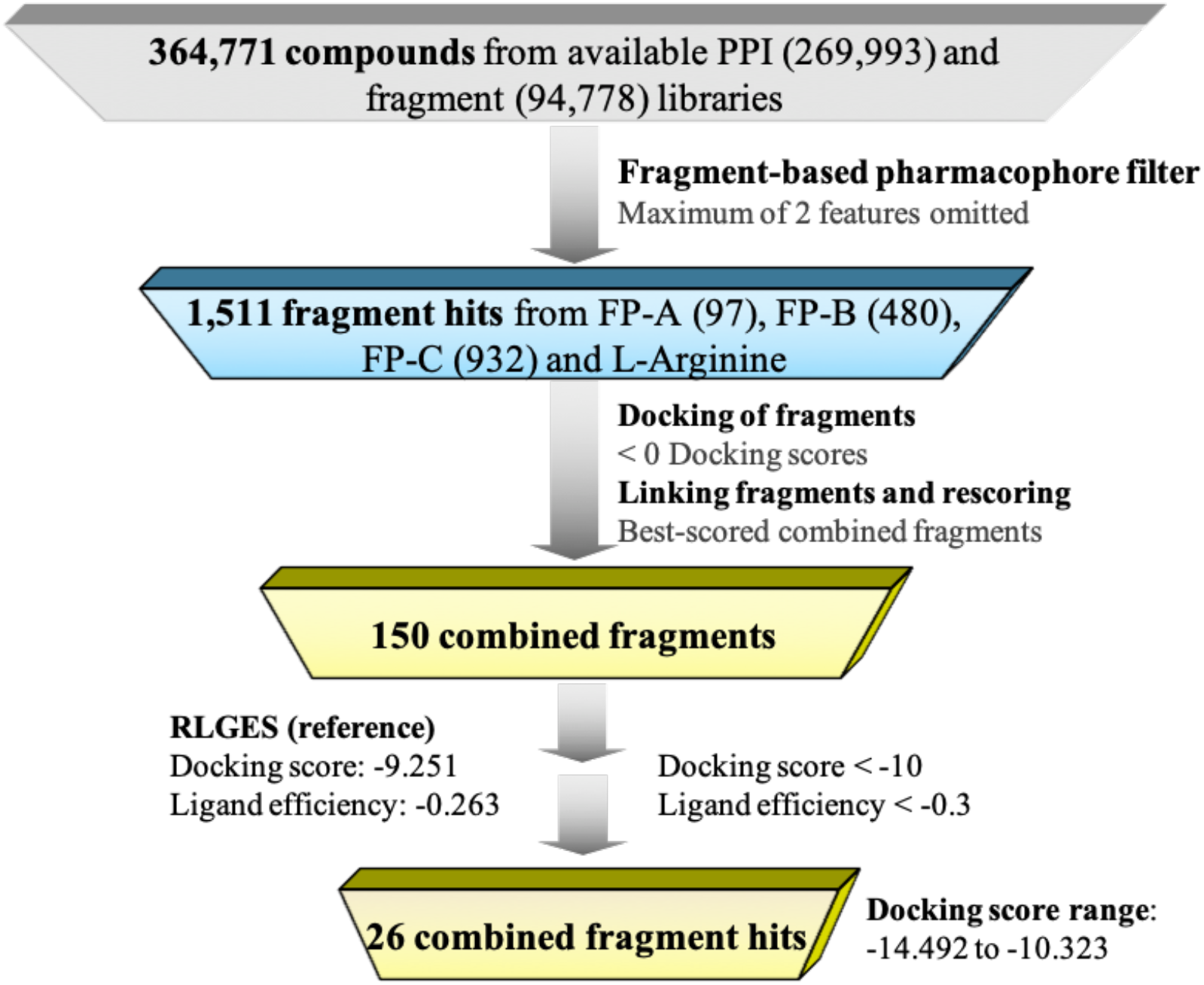
Overall virtual screening (VS) workflow. The different metrics of the RLGES peptide used as a reference are shown on the left, while the thresholds employed are shown on the right.

### Safety and molecular properties of the resulting combined fragments

The designed combined fragments were generated through an *in silico* process where low-complexity compounds were joined through methyl or methylene linkers. Therefore, it is crucial to evaluate the safety and the synthetic accessibility of these compounds before considering their application as E3 ligase recruiters. To that end, we calculated the quantitative estimation of drug-likeness (QED),[30] the synthetic accessibility (SA) score,[31] and the presence of Pan Assay Interference Compounds (PAINS)[32] using RDKit python functions.[33] QED ranges between zero (all properties unfavorable) to 1 (all properties favorable). SA score ranges between 1 (easy to make) to 10 (very difficult to make) and ideally, the number of PAINS should be 0. We started evaluating the RLGES peptide, which as expected, presents a significantly low drug-likeness (QED: 0.04). However, it is easy to synthesize (SA score: 4.9) and does not contain any PAINS compounds. Subsequently, the evaluation of the 26 combined fragments reveals a substantial enhancement in drug-likeness compared to the RLGES peptide (QED: 0.09-0.48). These combined fragments are also feasible to synthesize (SA score: 4.33-6.67) and are free from PAINS compounds. These molecular metrics provide an estimation of the safety of the combined fragments but they can be also used to further filter the number of combined fragment candidates. Since molecules with QED > 0.35 are considered desirable and SA scores < 6 are considered easy to synthesize, we applied these threshold values to obtain a reduced set of combined fragments. This resulted in 4 best-scored combined fragments, named BCF 1-4, see **Figure S1**. As expected, their respective docking poses showed that they can establish interactions with key residues at the binding interface (I174, D176, D179, S172, T144, and D142) as well as multiple interactions with the protein surroundings of the UBR box (K536, Q572, T575, E568, and R567), see **Figure 5**. Interestingly, all the designed recruiters exhibit solvent exposure areas, which highlights its potential to be linked to target protein ligands without affecting their binding affinity. This manageable number of compounds is suitable for a more comprehensive computational evaluation and investigation of their potential as E3 ligase UBR1 recruiters.

**Figure 5.**
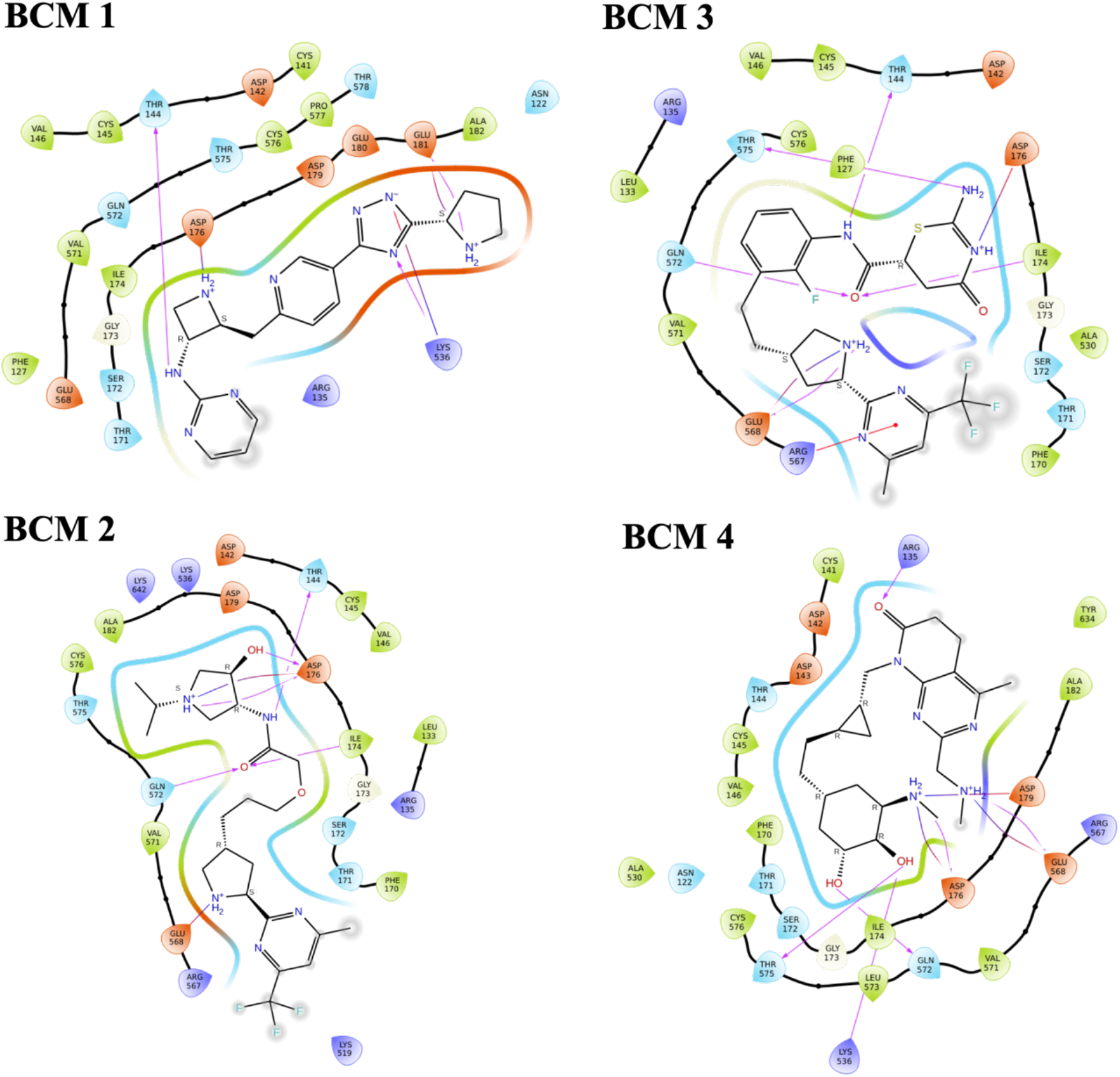
2D representation of the best-scored combined fragments interacting with UBR1. The UBR1 binding pocket residues are depicted as a function of the amino acid type: negatively charged are shown in red, positively charged in purple, polar in blue, and hydrophobic in green. Hydrogen bonds are drawn as pink arrows, pi-cation interactions as red lines with a dot, and salt bridges as a red-blue line. Atoms exposed to the solvent are indicated with a grey-shaded sphere.

### Evaluation of the binding free energy surface (BFES) of the best-scored combined fragments

The reconstruction of the BFES is essential to evaluate the performance of the designed recruiters. It informs about the relative stability of the recruiter in the E3 binding site and provides insights into the binding pathway and mechanism. Nonetheless, the reconstruction of the BFES is often challenging due to the long timescale of the binding process, requiring the use of advanced molecular dynamics simulation techniques.[34] To provide a preliminary assessment of the bound-state stability, we conducted conventional molecular dynamics simulations of the RLGES peptide and the BCF 1-4 compounds, using the docking poses as starting structures. These MD simulations showed that all compounds remain stable in the binding site of UBR1 for 100 ns (**Figure S2**). Given that, all the compounds were considered for the reconstruction of the binding pathway. To that end, we employed funnel metadynamics (FM) simulation, a recently enhanced sampling technique that has demonstrated great efficiency in computing the BFES for a variety of protein-ligand systems.[35-38] FM accelerates the BFES reconstruction by applying a bias potential that prevents the trapping of the system in a stable protein-bound conformation by applying a funnel-shaped restrain potential to restrict the ligand exploration to the region of interest, avoiding the sampling of the ligand around the entire bulk solvent. The funnel-shaped restrain consists of a conic region that encompasses the protein binding site connected to a cylindric region directed towards the solvent (see more details in **Figure S3**).

We started by computing the BFES of the RLGES wild-type peptide (**Figure 6** and **Figures S4-5**). The BFES of the RLGES peptide showed that the bound state is highly stable (**Figure 6**). As observed experimentally, the bound conformations exhibit a robust hydrogen bond network. This network encompasses water-mediated hydrogen bonds connecting the guanidine group of RLGES with residues D179, D176, D142, and T144, in addition to hydrogen bonds formed between the NH3+ group of RLGES and residues I174, D176, and Q572 (**Figure 6A**). Interestingly, we observe that the guanidine group can rotate and sample different poses, which leads to a dynamic water-mediated hydrogen bond network. At the C-terminal extreme of RLGES, the Ser residue forms hydrogen bonds with residues R567 and S172 while the Glu residue remains solvent-exposed thus exhibiting higher flexibility. The transient break of these hydrogen bonds leads to increased flexibility in the C-terminal region, resulting in the destabilization of the bound state, and causing a displacement of the peptide from the binding site. This displacement provokes a rearrangement of the guanidine water-mediated interactions and the loss of key peptide-UBR1 hydrogen bonds, such as the NH3+ with I174, D176, and Q572 (**Figure 6B**). At this point, the peptide rapidly evolves towards total unbinding conformations. Upon unbinding, the peptide mostly samples folded conformations, where the guanidine group of Arg and the carboxylic acid group of Ser approach each other forming a stable salt bridge (**Figure 6C**).

**Figure 6.**
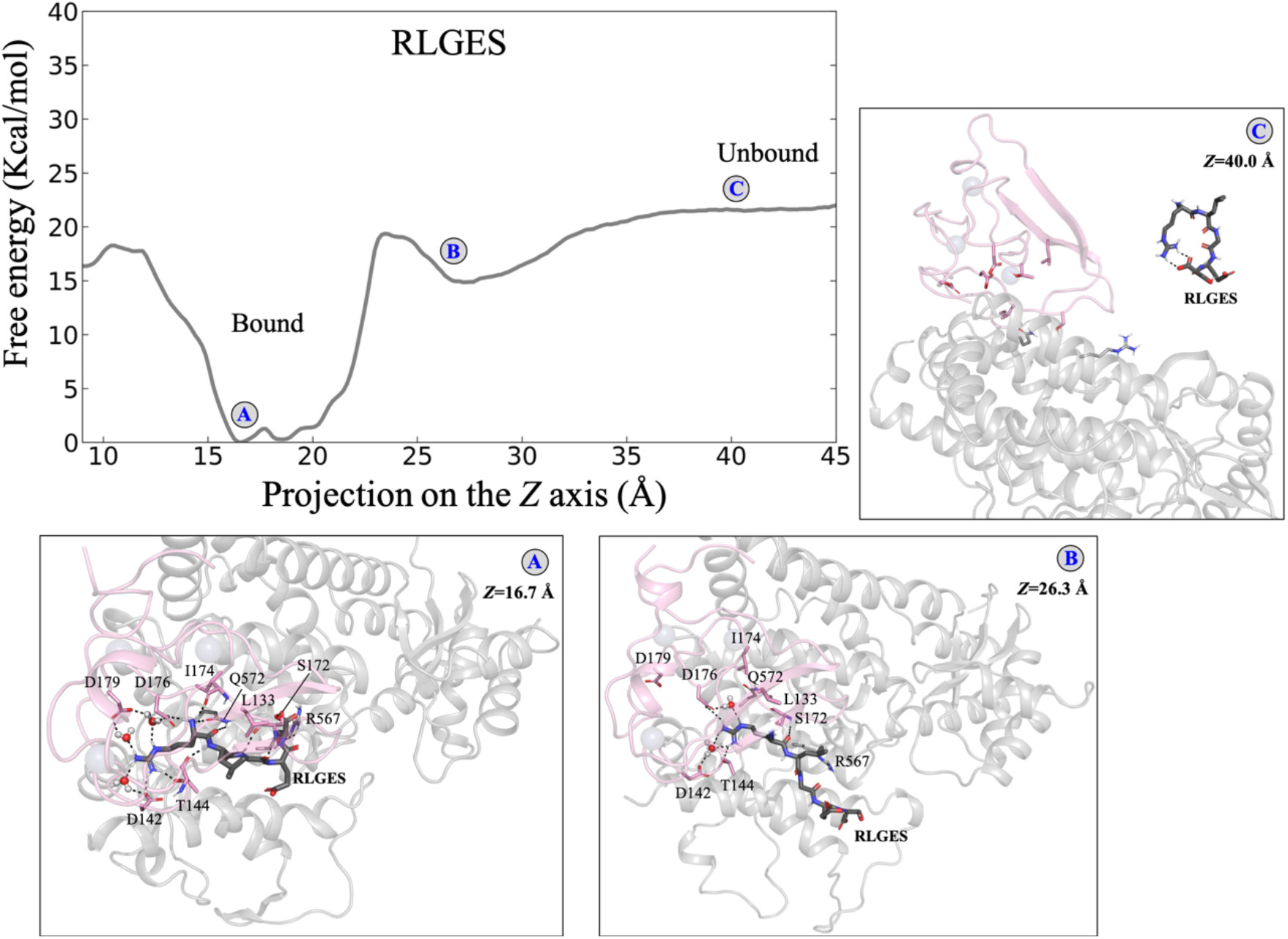
Binding Free Energy Surface (BFES) of the RLGES peptide. Reconstruction of the BFES as a function of the progression of the COM of the peptide along the Z axis of the funnel obtained from FM simulations. Representative conformations of the binding pathways (A, B, and C) are shown. The UBR box is colored pink while the protein surroundings in grey. The RLGES peptide is shown as dark grey sticks. The polar interactions between the RLGES peptide, water molecules, and protein residues are represented as black dashed lines.

The BFES profile of the RLGES wild-type peptide serves as a reference to evaluate the designed compounds. Considering the induced-degradation mode of action of PROTACs, compounds presenting BFES profiles with much lower binding free energy than RLGES may result in recruiters that bind efficiently to the UBR1 protein but do not properly dissociate to be available for the next target-protein degradation cycle. On the contrary, compounds presenting BFES profiles with much higher binding free energies may not efficiently bind to the UBR1 protein and therefore affect the degradation process of the target protein. The BFES profiles of the designed compounds reveal their potential as recruiters (**Figure 7-8** and **Figure S4-5**). BCF 1 presents the lowest free energy binding and thus the most stable binding pose. This can be attributed to a deeper inclusion into the binding cavity. As predicted by the docking pose, BCF 1 forms polar contacts with E181, D176, T144, and R536. However, the FM simulations reveal additional water-mediated contacts with D179 and I174, further stabilizing BCF 1 in the binding site. Upon unbinding, the pyrimidine and the pyridine rings perform pi-pi stacking interactions resulting in stable folded conformations. The significantly lower binding free energy of BCF 1 to RLGES (ΔΔG ca. -7.4 kcal/mol) suggests that the unbinding process may be too slow and infrequent to serve as an efficient recruiter. In contrast, BCF 2 showed a similar profile to that of the RLGES peptide, with a marginal ΔΔG increase of ca. 2.2 kcal/mol (**Figures 7-8**). BCF 2 is mainly anchored in the active site through polar interactions with D176, I174, Q172, T144, and E568 which can also involve water-mediated interactions (**Figure 7**). Remarkably, the trifluoromethyl group can be accommodated in the interface between the UBR box and protein surroundings, which can also substantially contribute to the binding energy. This also suggests that the methylene group, which is more solvent-exposed, is a better choice to link protein degradation ligands. Upon unbinding BCF 2 does not stabilize in folded conformations. For all these reasons, BCF 2 presents a suitable scaffold for recruiting E3 ligase UBR1. BCF 3 compound shares the fragment containing the trifluoromethyl group with BCF 2. Nevertheless, the simulations indicate that it anchors less effectively through the other fragment, resulting in a ΔΔG ca. 5.9 kcal/mol higher than BCF 2 (**Figure 8**). Lastly, the BFES for BCF 4 presents a very low binding free energy, suggesting that it is expected to be an inefficient recruiter.

**Figure 7.**
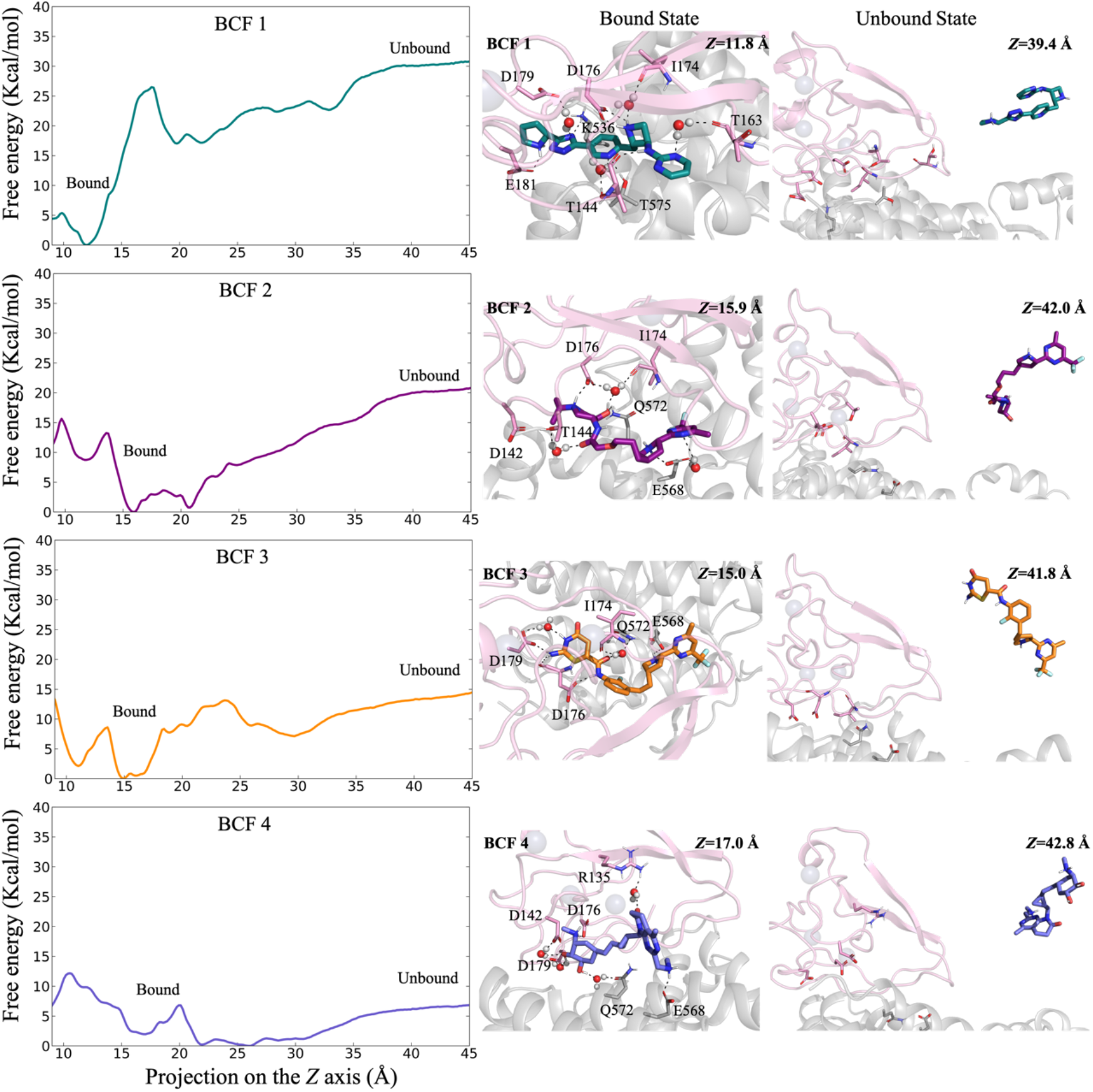
Binding Free Energy Surface (BFES) of BCF 1-4. Reconstruction of the BFES as a function of the progression of the COM of BCF1-4 along the Z axis of the funnel obtained from FM simulations Representative conformations of the bound and unbound states are shown. The UBR box is colored pink while the protein surroundings are grey. The BCF 1-4 molecules and their interacting residues are shown as dark grey sticks. The polar interactions between the BCF 1-4, water molecules, and protein residues are represented as black dashed lines.

**Figure 8.**
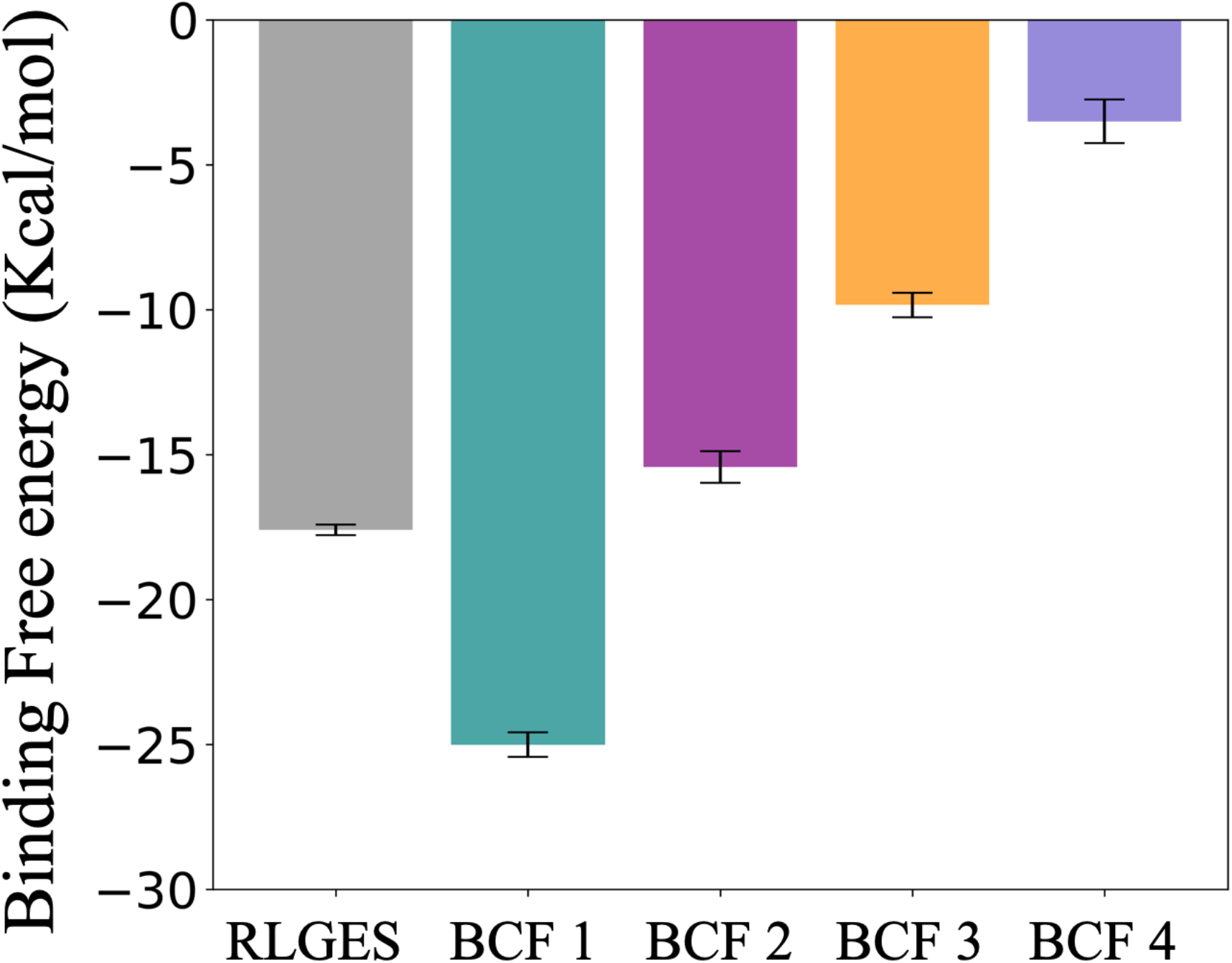
Binding Free Energy values of the RLGES peptide and BCF 1-4. These values were calculated as the average over the last 200 ns of the simulation with their corresponding uncertainties estimated as the standard deviation from these average values. Notably, the binding free energy values take into account an entropy correction of ca. 3.3 kcal/mol due to the utilization of the funnel shape potential in the unbound state. The entropy correction was calculated using the equations described in Ref. 35.

## CONCLUSIONS

Proteolysis targeting chimeric molecules (PROTACs) holds promising potential in drug discovery by facilitating specific degradation of disease-related proteins within cells. However, the clinical progression of PROTACs has faced significant obstacles, mainly attributed to their peptide-based structure and the scarcity of appropriate E3 ligases. Here, we concentrate on creating agents for recruiting the E3 ligase UBR1. This ubiquitously expressed protein has been recently shown to be highly effective in the degradation of oncogenic proteins. Through a fragment-based virtual screening strategy, we have identified fragments that exhibit characteristics resembling the various zones of the wild-type RLGES peptide when it is bound to the E3 ligase UBR1. Subsequently, an optimization process involving fragment-linking led to the creation of combined fragments. To verify their suitability as therapeutics, we assessed their safety and synthetic accessibility. These combined fragments are expected to mimic the physicochemical properties of the wild-type peptide while improving the disadvantages of the peptide such as stability towards proteolysis and transport properties. The reconstruction of the binding free energy surface (BFES) of the best-scored combined fragments showed their potential utility as recruiters. BCF 2 exhibits a similar BFES profile to that of the wild-type recruiter RLGES, making it the most promising design. The BFES calculations also revealed the binding mechanism of both the RLGES peptide and designed compounds. In all cases, we observed that a dynamic network of water-mediated hydrogen bonds was crucial for stabilizing the bound state. The collection of this dynamic conformational ensemble proves to be valuable for facilitating structure-based hit-to-lead procedures for the designed compounds. Furthermore, upon closer examination of these structures, it is evidenced that the designed E3 ligase UBR1 recruiter, BCF 2, can easily be linked to protein target ligands via its solvent-exposed methylene group, thereby enabling the development of efficient PROTACs. Utilizing the E3 ligase UBR1 as degradation machinery represents a novel approach that holds promise for degrading disease-relevant proteins independent of the cell type. Moreover, the fragment-based peptidomimetics strategy devised in this study can also be employed for modeling related PROTACs that entail intricate protein-protein interactions.

## MATERIALS AND METHODS

### Pharmacophore model and virtual screening

The pharmacophore model was generated through the LigandScout 4.4 [39] using the PDB 3NIN crystal structure, which contains the UBR box domain of *Saccharomyces cerevisiae* UBR1 in complex with the N-Degron peptide (RLGES) from cohesion subunit Scc1. For simplicity, two polar contacts with water molecules were manually removed. LigandScout uses a pattern recognition-based approach. First, all molecules from compound libraries are transformed into pharmacophores and second, it screens with pattern recognition the library of pharmacophores towards the given pharmacophore model. The results are the best matching pairs for each feature. For all fragment pharmacophore models (FP-A, FP-B, and FP-C) a maximum of two features were omitted during the screening process to increase the hit rates. The commercial fragment libraries screened are Life Chemicals (https://lifechemicals.com/screening-libraries/fragment-libraries), Asinex (https://www.asinex.com/building-blocks/fragments), Otava (https://otavachemicals.com/products/fragment-libraries/general-fragment-library), and ChemDiv (https://www.chemdiv.com/catalog/complete-list-of-compounds-libraries/).

### Docking virtual screening and fragment-linking optimization

The crystal structures of both the UBR box domain of *Saccharomyces cerevisiae* UBR1 (PDB 3NIN) and the whole UBR1 length (i.e. UBR box domain + protein surroundings) (PDB 7MEX) were prepared using the protein preparation Wizard module[40] [41] in Schrodinger Suite. It includes filling in missing side chains through Prime,[40] the assignment of protonation states and H-bond optimization through PROPKA, and an all-protein atom minimization using the OPLS4 force field.[42] The 1,511 fragment hits obtained from the pharmacophore virtual screening were prepared using the LigPrep module[43] in Schrodinger Suite, where for each fragment, all ionizable states significantly populated at pH 7.0 +/-2.0 were generated through Epik,[44] and all structures were energy minimized using the OPLS4 force field.[42] For the docking calculation, we used the Ligand Docking Glide XP (Extra-Precision) module[45] of Schrodinger Suite. As recommended by Schrodinger for fragment docking, we increased the initial number of poses per ligand from 5,000 to 50,000; widened the scoring window from 100 to 500; increased the number of minimized poses per ligand to 1,000; and used the expanded sampling. Subsequently, we employed the “Combine Fragments” module in the Schrodinger Suite to covalently link the pre-positioned fragments on the UBR1 binding site. This method identifies bonds that can be formed between fragments according to geometric criteria. To ensure a proper rotational alignment of the fragments, the angle between the bond directions was set to be less than 15°. To ensure a proper translational alignment of the fragments, the distance between the atom in one fragment and the atom in the other fragment was set to be less than 1 Å. For fragments that have no atoms close enough to create new bonds but are still not far apart, a methylene linker per fragment can be included to try to link those pairs of fragments. Fragments were joined in one single round, i.e., combined fragments were no further subjected to join a third fragment. Finally, the resulting combined fragments were ranked and only the top 150 structures were retrieved.

### Molecular dynamics simulations

*System preparation:* The E3 ubiquitin-protein ligase UBR1 (PDB 7MEX) presents a length of 1,950 residues. Given the extensive size of the protein, we constructed a system that comprises the UBR box domain (82 residues) and its protein surroundings (419 residues), see **Figure S3**. Thus, we manually cut the protein chains and added acetyl (Ace) and *N*-methylamine (Nme) caps in the C and N-terminal residues, respectively. In total, we constructed 5 systems, the UBR1 in complex with the RLGES wild-type peptide, and the best scored combined fragments (BCF 1-4). For all ligands, the staring pose was derived from the docking calculations. MD parameters for the combined fragments (BCF 1-4) were generated with the parmchk module of AMBER20[46] using the general amber force field (GAFF). The atomic charges were obtained using the AM1-BCC model[47] through the antechamber module of AMBER20.[46] All systems were filled into a pre-equilibrated cubic box with a 25-Angstrom buffer of water molecules and neutralized by the addition of explicit counterions (Na^+^ and Cl^-^). *Molecular dynamics protocol:* The systems were minimized in a two-stage geometry optimization approach. First, a short minimization of the water molecules and counterions positions, with positional restraints on the protein and ligands was performed at constant volume periodic boundary conditions. Second, an unrestrained minimization including all atoms in the simulation cell was carried out. All systems were heated using seven 50 ps steps, incrementing the temperature ca. 50 K each step (0-298.15 K). Decreasing harmonic restraints were applied to the protein (210, 165, 125, 85, 45, and 10 Kcal/mol Å^2^) during the thermal equilibration, with the Langevin thermostat used to equalize and control the temperature. During the heating process, the initial velocities were randomized. For the heating and following steps, bonds involving hydrogen were constrained with the SHAKE algorithm and the time step was set to 2 fs, allowing potential inhomogeneities to self-adjust. An 11 Å cutoff value was applied to Lennard-Jones and electrostatic interactions. The equilibration step was performed in three stages. In the first stage, an MD simulation of 1 ns under NVT ensemble and periodic boundary conditions was performed to relax the simulation temperature. In the second stage, an MD simulation of 2 ns under an NPT ensemble at a simulation pressure of 1.0 bar was performed to relax the density of the system. In the third stage, an MD simulation of 2 ns under an NVT ensemble was performed to further relax the density/volume defined by pressure. After the equilibration of the system, we performed an MD production run of 100 ns for each system.

### Funnel metadynamics (FM) simulations

FM is a powerful method to reconstruct the binding free energy surface of protein-ligand complexes as a function of a few user-defined descriptors, also referred to as collective variables (CVs).[34] In this work, we selected the progression of the center of mass (COM) of the ligand along the Z-axis of the funnel (see **Figure S3** for visualization and customization parameters of the funnel shape potential). The simulations were carried out using the Amber20 code[46] together with the Plumed-master plugin, which contains the FM module v2.0.[38, 48] During an FM simulation, external energy quantities (Gaussian functions) are added to the CV values along the simulation time. This external potential encourages the system to escape from local energy minima and overcome energy barriers, thus allowing for enhanced sampling of the CV space. After sufficient simulation time, the bias potential converges and the FEL can be reconstructed by summing the Gaussian functions added to the CV values along the simulation time. Here, Gaussian functions of height 2.0 kJ mol-1 and width 0.01 were deposited every 2 ps of the MD simulation at 300 K. For a smooth convergence of the bias potential, we used the well-tempered (WT) version of the metadynamics algorithm, in which the height of the gaussian potentials were gradually decreased over time proportional to the potential deposited in the currently visited point of the CV space. A bias factor parameter of 20 was selected to control how quickly the Gaussian height is decreased. In addition, the multiple-walkers approach was used to improve the conformational sampling and to speed up the metadynamics simulations. It is based on running in parallel interacting replicas (walkers) where each walker biases the identical CV and reads the Gaussian functions deposited by the others during the simulation, thus reconstructing the same metadynamics bias simultaneously. In particular, we run 5 walker replicas (W1-5) for each system. The structures used as a starting point were the ligand-bound docking poses after system minimization and equilibration as described in the molecular dynamics protocol. After 1-1.2 μs of accumulated simulation time for each system, we reconstructed their binding free energy surfaces by summing the Gaussian potentials deposited by all walker replicas as a function of the CV (see **Figures S4-5** for convergence assessment).

## ACKNOWLEDGEMENTS

This work was supported by the Bio & Medical Technology Development Program (NRF-2022M3E5F3080873), the Mid-career Researcher Program (NRF-2020R1A2C2101636), the Medical Research Center (MRC) grant (NRF-2018R1A5A2025286), and the Brain Pool Program (NRF-2021H1D3A2A02038434) funded by the Ministry of Science and ICT (MSIT) through the National Research Foundation of Korea (NRF). It was also supported by the Ewha Womans University Research Grant of 2021. We are grateful to the Korea Institute of Science and Technology Information (KISTI) Supercomputing Center for providing computing resources and technical assistance (KSC-2022-CRE-0517). We thank Prof. Hyun-Suk Lim for the helpful discussions and comments on the manuscript.

