## Supplementary Information for "IN SILICO DESIGN AND BINDING MECHANISM OF E3 LIGASE UBR1 RECRUITERS"

| Commercial Libraries | FP-B hits | FP-C hits | FP-A hits |
|---|---|---|---|
| **PPI**<br>(ippdb, Asinex, ChemDiv, Otava and Life Chemicals) | **173**/269,993<br>(0.06 %) | Not Tested | **30**/269,993<br>(0.01 %) |
| **Fragment**<br>(Asinex, ChemDiv, Otava and Life Chemicals) | **340**/94,778<br>(0.4 %) | **967**/94,778<br>(1 %) | **72**/94,778<br>(0.08 %) |
| **Total**<br>(removing duplicates) | **480**/ 364,771<br>(0.1 %) | **932**/94,778<br>(1 %) | **97**/ 364,771<br>(0.03 %) |

**Supplementary Table 1**. Summary results of the fragment-based pharmacophore virtual screening. The virtual hit numbers are highlighted in bold and the hit rates are shown in brackets. In all screening process, a maximum of two pharmacophore features could be omitted.

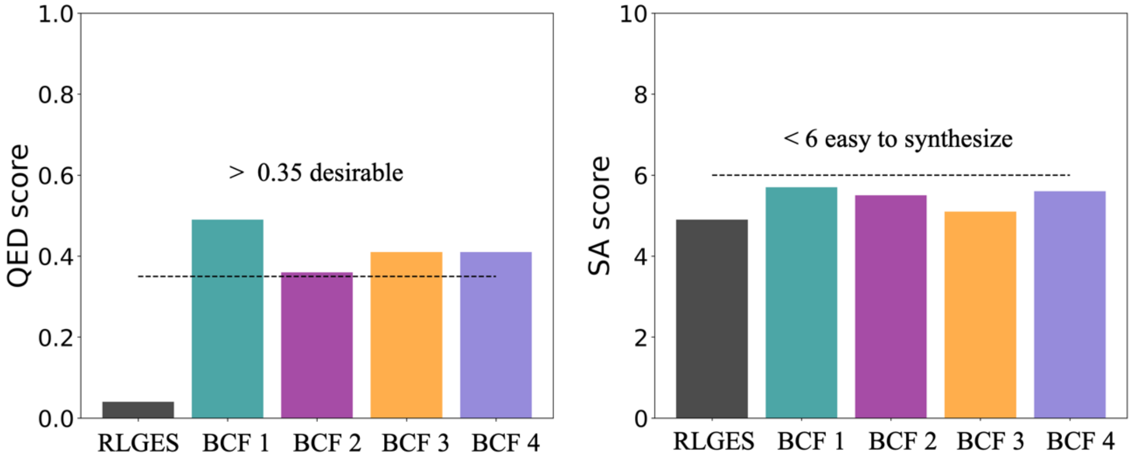


**Figure S1**. **Drug-likeness and synthetic accessibility of the N-Degron peptide (RLGES) and designed BCF 1-4**. The Quantitative Estimation of Drug-likeness (QED) and Synthetic Accessibility (SA) score thresholds are depicted by a dashed line.

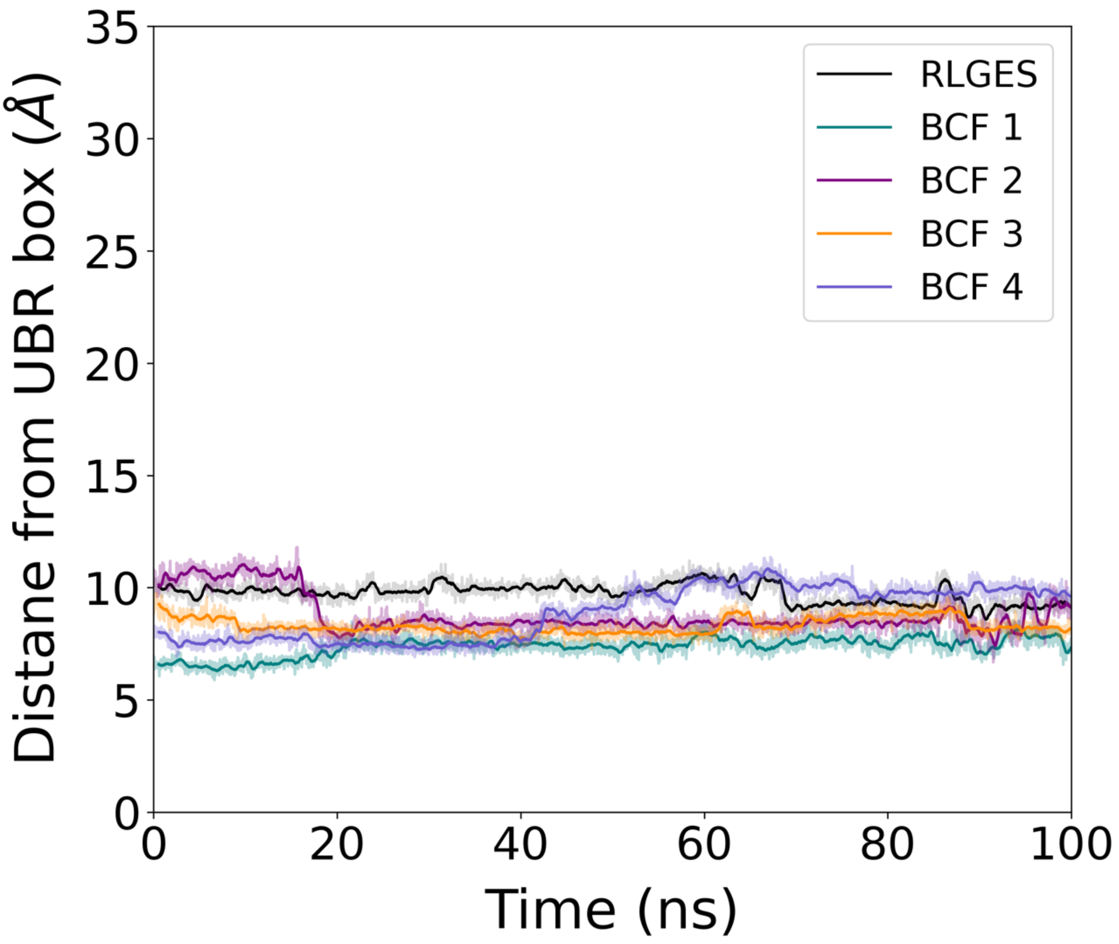


**Figure S2**. **Stability analysis of the ligands in the UBR 1 binding site obtained from molecular dynamics simulations.** The distance from the UBR box corresponds to the distance between the COM of the ligands and the COM of the I174 backbone (located at the UBR box binding site).

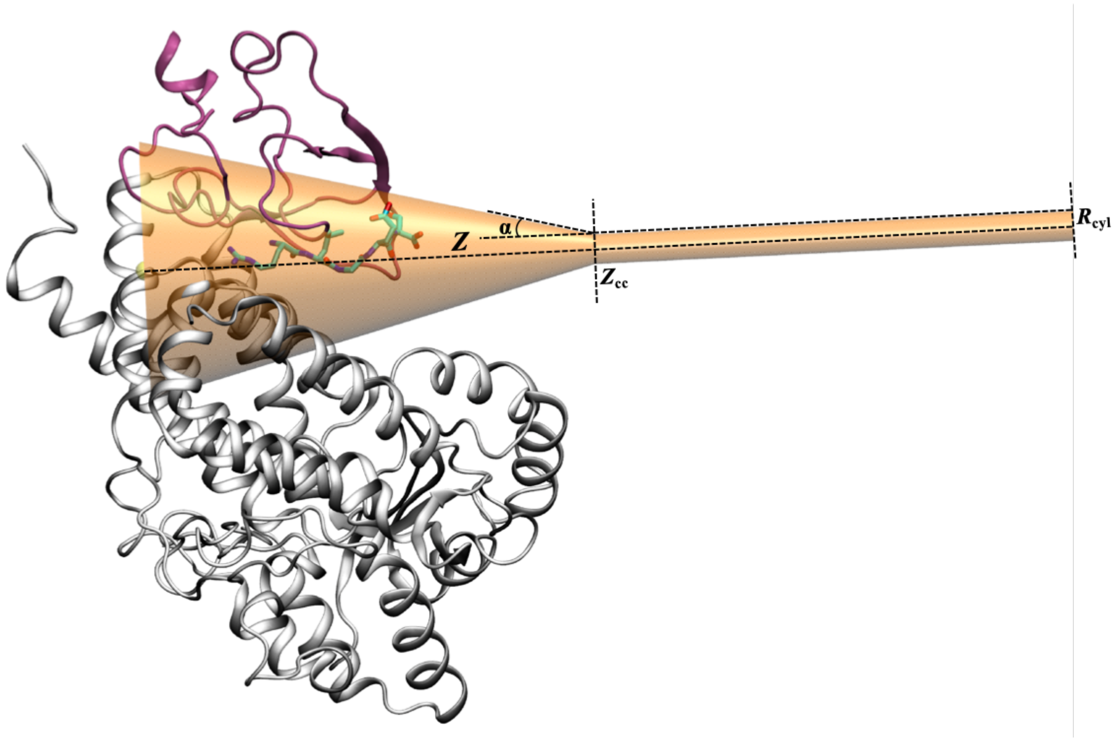


**Figure S3**. **Representation of the UBR1 protein region and the funnel-shaped restrain potential used in this work.** The UBR1 box domain (82 residues, in pink) and the UBR1 protein surroundings (419 residues, in grey) are shown in cartoon style while the RLGES wild-type peptide is in cyan sticks. The conic and cylindrical regions of the funnel restrain potential are depicted in orange. *Z* defines the axis the COM of the ligands progresses along the binding pathway. The funnel shape was customized as follows: the distance where the funnel shape switches from a cone into a cylinder ($Z_{cc}$) was set to 45 Å, the angle defining the amplitude of the cone (α) to 0.25 rad, and the radius of the cylindric region ($Z_{cyl}$) to 1.5 Å.

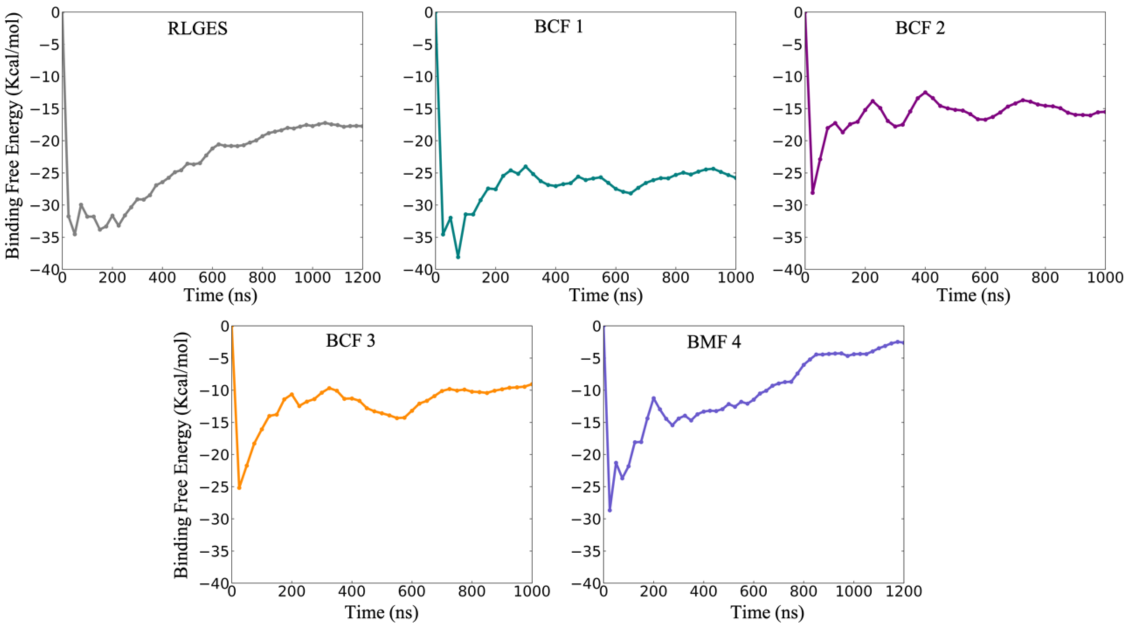


**Figure S4**. **Convergence of the binding free energy for the N-Degron peptide (RLGES) and the designed BCF 1-4**. The lines represent the free energy differences between the bound and bound state regions along the simulation time. The binding free energies were calculated every 25 ns. In the last part of the simulation, the binding free energies do not change significantly, which indicates convergence. Accordingly, the binding free energy values used in this work were calculated as the average over the last 200 ns of the simulation, with their corresponding uncertainties estimated as the standard deviation from these average values. The binding free energy values take into account an entropy correction of ca. 3.3 kcal/mol due to the utilization of the funnel shape potential in the unbound state. The entropy correction was calculated using the equations described in Ref. 35.

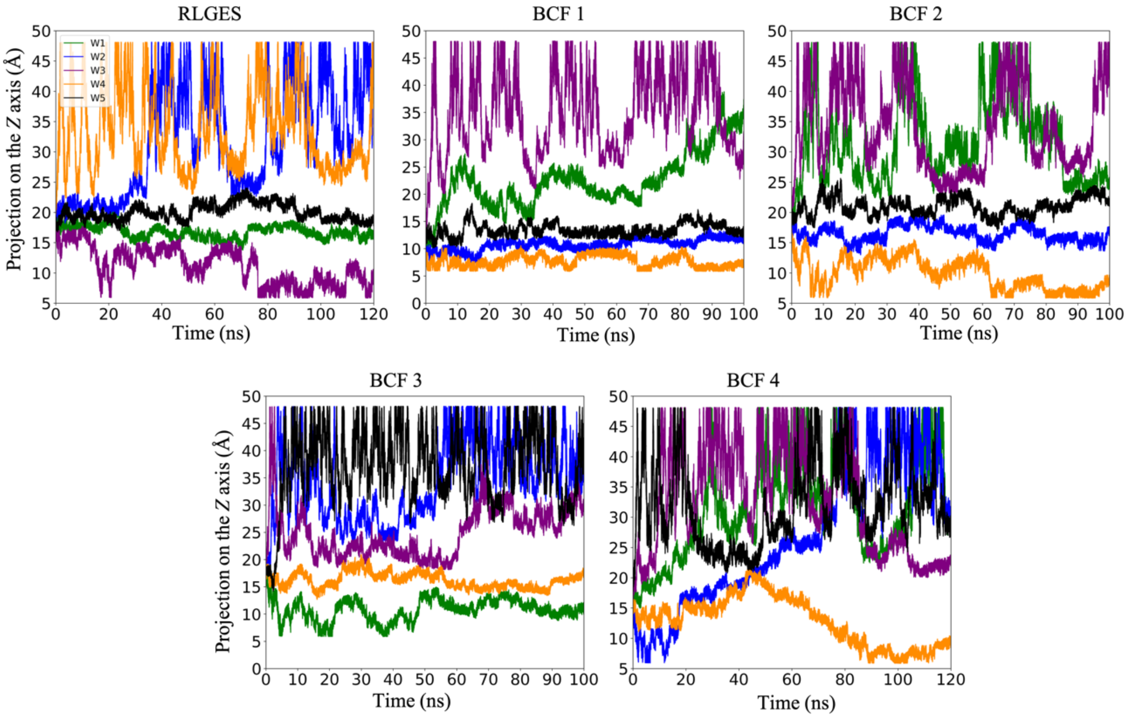


**Figure S5**. **Evolution of the COM of the BCF 1-4 on the Z axis of the funnel shape over the simulation time.** All walker replicas (W1-5) were started from the bound state in all systems. The plots show that the bias potential applied during the FM simulations encourages the walkers to sample the binding pathway and explore re-crossing events between bound and unbound states, thereby successfully reconstructing the binding free energy surface.
